# Early functional and cognitive declines measured by auditory evoked cortical potentials in Alzheimer’s disease mice

**DOI:** 10.1101/2021.02.25.432945

**Authors:** Ling Mei, Li-Man Liu, Kaitian Chen, Hong-Bo Zhao

## Abstract

Alzheimer’s disease (AD) is characterized with a progressive loss of memory and cognitive decline. However, assessment of AD-associated functional and cognitive changes is still a big challenge. Auditory evoked cortical potential (AECP) is an event-related potential reflecting not only neural activation in the auditory cortex but also cognitive activity in the brain. In this study, we used the subdermal needle electrodes with the same electrode setting as the auditory brainstem response (ABR) recording and recorded AECP in normal aging CBA/CaJ mice and APP/PS1 AD mice. AECP in mice usually appeared three positive peaks, i.e., P1, P2, and P3, and three corresponding negative peaks N1, N2, and N3. In normal aging CBA mice, the early sensory peaks P1, N1, and P2 were reduced as age increased, whereas the later cognitive peaks N2, P3, and N3 were increased or had no changes with aging. Moreover, the latency of P1 peak was increased as age increased, although the latencies of later peaks had significant reduction with aging. In AD mice, peak P1 was significantly reduced in comparison with wild-type (WT) littermates at young ages, proceeding AD phenotype presentation. In particular, the later cognitive peak P3 was diminished after 3 months old, different from normal aging effect. However, the latencies of AECP peaks in AD mice generally had no significant delay or changes with aging. Finally, consistent with AECP changes, the accumulation of amyloid precursor protein (APP) at the auditory cortex was visible in AD mice as early as 2 months old. These data suggest that AECP could serve as an early, noninvasive, and objective biomarker for detecting AD and AD-related dementia.

## Introduction

Alzheimer’s disease (AD) is a progressive neurodegenerative disease, characterized by a progressive loss of memory and dementia. Over the last two decades, the prevalence of AD and AD-related dementia (ADRD) has been rapidly increased. It is prospected that there will be 150 million AD patients by the year 2050, almost triple the population in 2018 (Hebert et al., 2013; Patterson et al., 2018). This will produce a heavy economic and social burden. Early detection is critical for prevention and interventions of this common neurodegenerative disease. It is estimated that early diagnosis and interventions delaying the onset of dementia by even one year could decrease the worldwide prevalence of dementia by 10% (Patterson et al., 2018). However, since lack of the early, reliable AD biomarkers, early detection of AD currently is still a big challenge. In particular, it lacks objective biomarkers to detect cognitive decline, which is the most common and predominant symptom of AD, at the early stage.

Recently, increasing evidence suggests that defects in sensory systems, including the olfactory, visual, and auditory systems, are highly associated with AD progressing (Murphy, 2019; Rochoy et al., 2019). AD pathology could appear in sensory associated areas before appearing in regions involving memory, such as the entorhinal and hippocampal areas (Gates et al., 2002; Murphy, 2019). These neuropathological changes in the sensory associated areas could also occur before cognitive symptoms become apparent. Hearing is an important neural sense. AD can cause hearing loss, and hearing loss also can in turn accelerate AD and dementia development (Morrison et al., 2018; Griffiths et al., 2020). Our previous study demonstrated that auditory brainstem response (ABR), which reflects the function and integrity of the auditory pathway in the brainstem, had reduction in APP/PS1 AD mice at the age of 3 months old (Liu et al., 2020), prior to occurrence of typical AD phenotypes, such as spatial learning deficit, in this AD mouse model by age of 6-7 months old (Reiserer et al., 2007; Ordonez-Gutierrez et al., 2015, 2016). Recent studies also found that deficits in auditory gap detection could occur in young 5xFAD mice (Kaylegian et al., 2019; Weible et al., 2020). These studies suggest that AD could cause hearing function decline at the early stage, and that hearing function tests could provide important information for AD development at the early stage.

Auditory evoked cortex potential (AECP) is an event-related potential evoked by auditory stimulation and reflects the neural activity in the auditory cortex and the related brain areas. In humans, AECP consists of several waveforms and are named as P1 (P50), N1 (N100), P2 (P200), N2 (N250), P3 (P300), and P4 (P600) based on their latencies (Swords et al., 2018). Early waveforms (i.e., P1 & P2) are “sensory”, largely depending on the stimulus and activities in the auditory cortex; later waveforms (i.e., P3 & P4) are “cognitive” reflecting cognitive processing (Sur and Sinha, 2009). Thus, unlike ABR, the AECP can reflect not only the neural activities in the auditory cortex and also cognitive activities in the brain.

In previous studies, auditory evoked potentials were recorded in the cortex, hippocampus, and other brain areas with implanted electrodes in AD animal models (Wang et al., 2003; Stoiljkovic et al., 2019) or amyloid-β (Aβ) injected in the rat and mouse brain at different ages (Kim et al., 2020; Hidisoglu and Yargicoglu, 2020). In general, auditory evoked potentials in the auditory cortex and other brain cortexes were reduced in AD or amyloid injected animals. However, there were conflicting results in the literature (Gurevicius et al., 2013). AD-induced auditory evoked potential changes in the brain and their dynamic changes in AD development still remain largely undetermined. In this study, we systematically recorded AECP in APP/PS1 AD mice to assess AD-induced functional changes in the auditory cortex and brain with aging. We found that AECP in AD mice had significant reduction at young ages. In particular, later cognitive peak P3 in AD mice was diminished, different from normal aging effect. Our study suggests that AECP recording could serve as an early noninvasive biomarker to assess AD and ADRD development.

## Materials and Methods

### APP/PS1 AD mice and CBA/CaJ normal aging mice

APP/PS1 transgenic mice were purchased from Jackson Laboratory, USA (Stock No: 004462, mixed C57BL/6;C3H genetic background). Since the C57BL/6 genetic background is associated with age-related hearing loss, we crossed APP/PS1 mice with CBA/CaJ mouse strain (Stock No: 00654, Jackson Laboratory, USA), which has no apparent age-related hearing loss (Zheng et al., 1999), for 4 generations. APP/PS1 transgenic mice possess both chimeric mouse/human amyloid precursor protein (Mo/HuAPP695swe) and mutant human presenilin 1 (PS1-dE9) (Reiserer et al., 2007). According to Jackson Laboratory’s technical notice, APP/PS1 transgenes were detected by PCR amplification of tail genomic DNA using the following primers: wild-type (WT) forward: 5’-GTG TGA TCC ATT CCA TCA GC-3’; Common: 5’-GGA TCT CTG AGG GGT CCA GT-3’; Mutant forward: 5’-ATG GTA GAG TAA GCG AGA ACA CG-3’. Mutant and WT mice generated 142 and 265 bps bands, respectively. WT littermates served as controls. Both genders were used in the experiment. CBA/CAJ mice were also used for normal aging study. All experiments and procedures were approved and conducted in accordance with the policies of the University of Kentucky Animal Care & Use Committee (approved protocol: UK#2020-3524).

### Auditory evoked cortical potential recording

Auditory evoked cortical potential (AECP) was recorded in a double-wall sound isolated chamber by use of a Tucker-Davis R3 system with an ES-1 high-frequency speaker (Tucker-Davis Tech. Alachua, FL). Mice were anesthetized by intraperitoneal injection with a mixture of ketamine and xylazine (8.5 ml saline+1 ml Ketamine+0.55 ml Xylazine, 0.1 ml/10 g). Body temperature was maintained at 37–38°C by placing anesthetized mice on an isothermal pad. As the same as ABR recording (Liu et al., 2020), two subdermal needle electrodes were punched into the head skin at the vertex and ventrolaterally to the right or left ear. The ground needle electrode was inserted to the right leg. Cortical potentials were evoked by clicks (85 dB SPL) in alternative polarity with stimulating rate of 1/s. The recording time window was 800 ms, starting from the beginning of the stimulation. The signal was amplified by 50,000 and averaged by 200 time. The recording filter was set at 0 to 3 kHz. The signal recording was performed by BioSigRP software (Tucker-Davis Tech). The collected signals were further filtered in off-line with a build-in 10-order digital Gaussian filter with 12.5 Hz of central frequency and a bandwidth of 2 Octaves for analysis.

### Brain slide preparation and immunofluorescent staining

Mice were anesthetized with Ketamine as described above and intracardially perfused with 4% paraformaldehyde for 15-20 min. After decapitation, the mouse skull was opened. The brain was carefully removed and incubated in 4% paraformaldehyde overnight for post-fixation. Following infiltration with 30% glucose for three day with changing fresh solution at each day, the brain was embedded in OCT (Cat # 4583, Sakura Finetek USA Inc. CA) at room temperature for one day. Then, the embedded brain was frozen at -80 °C overnight and cut 20 µm thick at -22∼24 °C by a cryostat (Thermo Electron Corp. Waltham, MA). The tissue sections were directly mounted onto glass slides for staining and storage.

Immunofluorescent staining was performed as described in our previous reports (Zhao and Yu, 2006; Wang et al., 2009). After 1 hr of incubation with the blocking solution (10% goat serum and 1% BSA in PBS) with 0.5% Triton X-100 at room temperature, the brain slides were incubated with primary antibody rabbit monoclonal anti-amyloid precursor protein (APP) [clone Y188] (#AB32136, Abcam, USA; 1:500) in blocking solution at 4 °C overnight. After washout with PBS three times, the brain slides were incubated with corresponding secondary antibody conjugated with Alexa Fluor-568 (Thermo Fisher Sci, USA) for 2 hr at room temperature. At the last 15 min, 4’, 6-diamidino-2-phenylindole (DAPI, 0.1 mg/ml, D1306; Molecular Probes) was added into the incubation solution to visualize cell nuclei. After completely washing out the second antibodies with PBS, the sections were mounted with a fluorescence mounting medium H-1000, Vector Lab, CA) and observed under a Nikon A1R confocal microscope system. The AD and WT mouse sections were simultaneously stained and observed with the same parameters.

### Data processing, statistical analysis, and reproducibility

The amplitude and latency of peaks in AECP were measured by a house-made program with MATLAB (Mathworks Inc. USA). Data were plotted by SigmaPlot and statistically analyzed by SPSS v25.0 (SPSS Inc. USA). Data were expressed as mean ± s.e.m. Student t tests were performed between APP and WT groups at the defined age. When three or more groups were compared, one-way ANOVA with Bonferroni post hoc test was used. The threshold for significance was α = 0.05. The AECP was recorded from mice generated by more than 15 different breeding pairs.

## Results

### Auditory evoked cortex potential (AECP) in mice

Fig. 1 shows AECP recorded from mice. The AECP recorded from mice usually appeared 3 positive peaks, i.e., P1, P2, and P3, and three corresponded negative peaks N1, N2, and N3 (Fig. 1). Like AECP recorded from humans, both early sensory peaks (P1, N1, and P2) and later cognitive waves (P3 and N3) were visible in the recording. However, the later cognitive wave P3 in mice was not clearly visible until the age of 3 months old (Fig. 1A&B). At 6 months old, AECP in mice was well-developed and P1, P2, and P3 peaks were clearly visible (Fig. 1C). The latencies of peaks P1, P2, and P3 were 13.5±1.89, 67.5±2.74, and 128.1±6.76 ms, respectively (Fig. 2A).

**Fig. 1.**
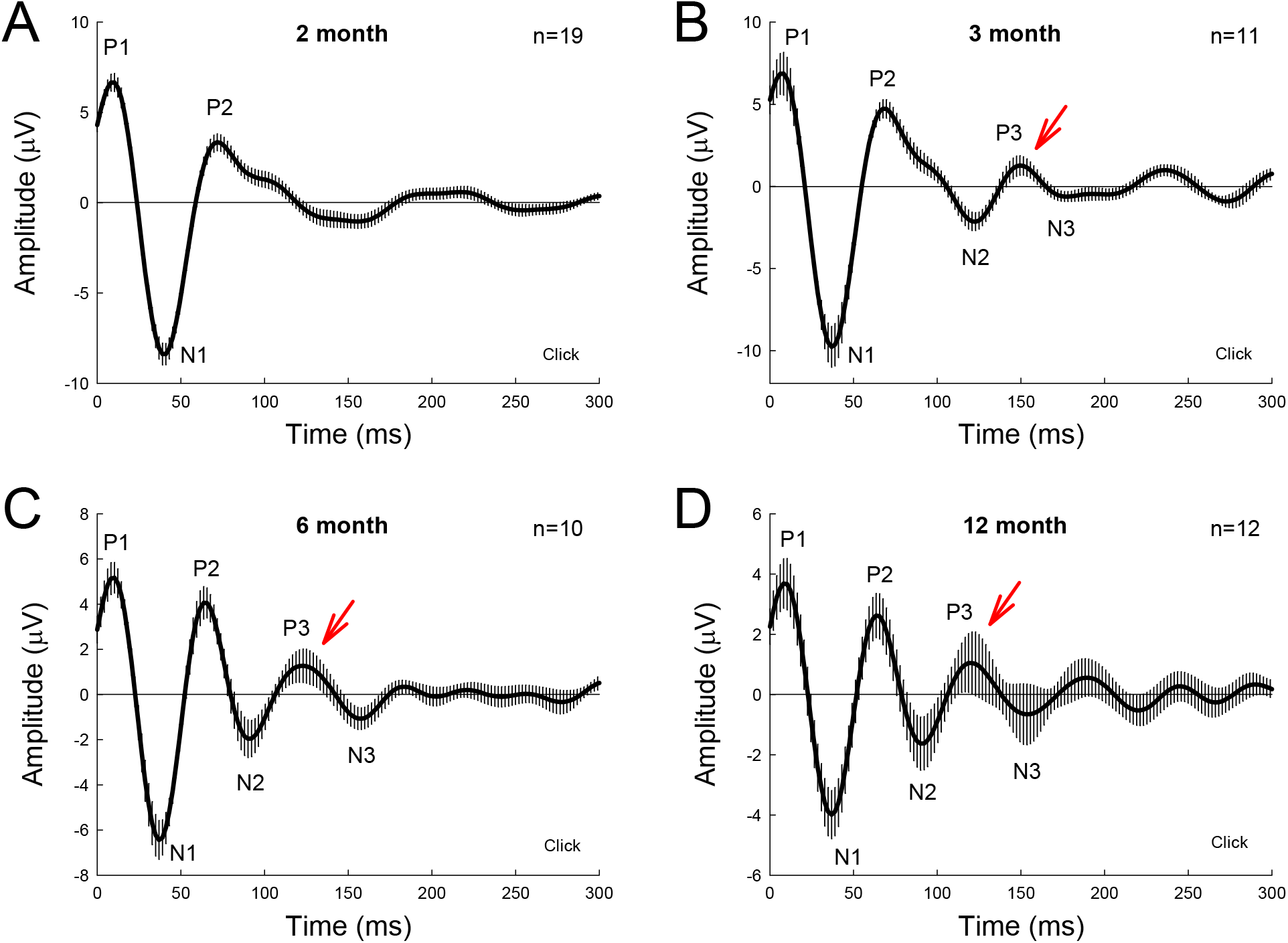
Auditory evoked cortex potential (AECP) in CBA/CaJ mice at different ages. AECP was evoked by a click stimulus (85 dB SPL) in alternative polarity. Traces were averaged from the recorded mice with both genders. Error bars represent SEM. Mouse age is indicated in each panel. N is the mouse number. AECP in mice has three positive peaks, i.e., P1, P2, and P3, and three corresponding negative N1, N2, and N3 peaks. Red arrows indicate P3 peak that is clearly visible in AECP at 3 months old and afterward.

**Fig. 2.**
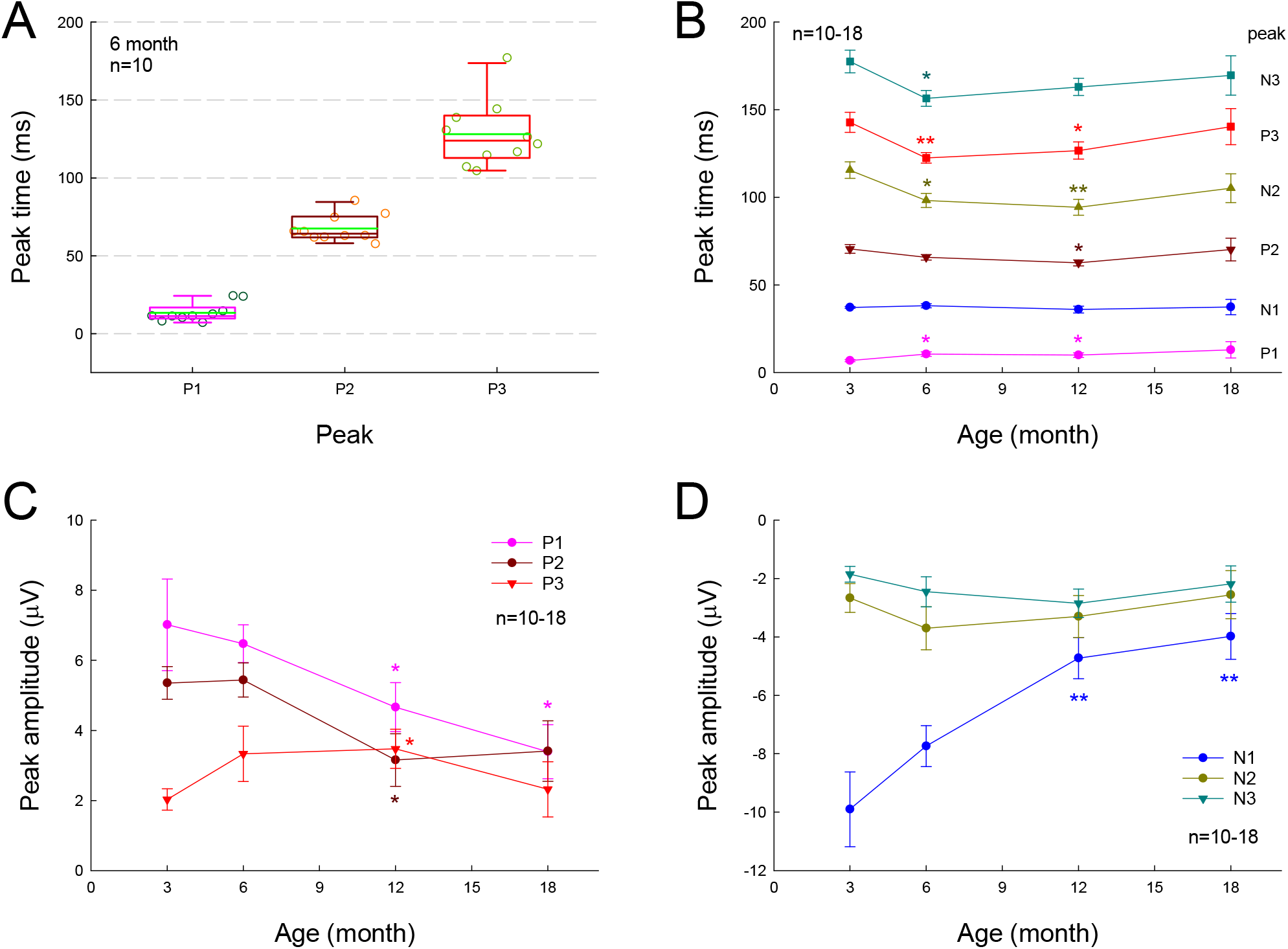
Changes of AECP in normal aging CBA/CaJ mice. N is the mouse number. **A:** P1, P2, and P3 peak-times at the age of 6 months old. The green line in the box represents the mean level. **B:** Changes of AECP peak-times in normal aging. The statistical t tests were performed by comparing peak-times at the defined age to that at 3 months old in each peak. **C-D**: Changes of peak-amplitudes with aging. The amplitudes of early peaks P1, N1, and P2 in AECP are decreased with aging, whereas the peak-amplitudes of later waveforms N2, P3, and N3 are increased or have no significant change with aging. The statistical analyses were performed by comparing peak-amplitudes at the defined age to that at age of 3 months old in each peak. *: P<0.05, **: P<0.01, t test, two-tail.

### Normal aging changes of AECP in CBA/CaJ mice

We further investigated the effect of normal aging on AECP. CBA/CAJ mice, which have no apparent age-related hearing loss (Zheng et al., 1999), were used. Fig. 1 shows that AECPs in CBA mice were reduced with age increased. The amplitudes of early peaks (i.e., P1, N1, and P2) were 4.66±0.70, -4.73±0.71, and 3.15±0.75 µV, respectively, at 12 months old and 3.39±0.77, -3.98±0.78, and 3.41±0.86 µV, respectively, at 18 months old (Fig. 2C&D). In comparison with those at the age of 3 months, the amplitudes of P1, N1, and P2 at 12 months old had significant reduction (P=0.003-0.022, t test, two-tail). However, the peak P3 was significantly increased from 2.03±0.31 µV at 3 months to 3.47±0.56 µV at 12 months old (P=0.037, t test, two-tail) (Fig. 2C). The amplitudes of later peaks N2 and N3 had no significant changes with aging (Fig. 2D).

The latency of P1 peak had significant increase at 6 and 12 months (P=0.014 and 0.045, respectively, t test, two-tail) (Fig. 2B). However, there was no significant change in the N1 latency with aging; the latencies of peaks P2, N2, P3, and N3 were significantly reduced (P=0.004-0.045, t test, two-tail) at 6 and 12 months old but had no significant changes at 18 months old in comparison with them at 3 months old (Fig. 2B).

### AECP in APP/PS1 AD mice

Fig. 3 shows AECP traces recorded from APP/PS1 AD mice and WT littermates at different ages. In general, P1 and P2 in AECP in AD mice appeared smaller than those in WT littermate mice at young ages. At the age of 1-3 months old, there was a notch at the descending side of peak P2 in AD mice (indicated by blue arrows in Fig. 3A-C). P3 is clearly visible in WT mice after 4 months old (indicated by red arrows in Fig. 3D-I), but small or invisible in AD mice (Fig. 3D-I). Later negative peaks of N2 and N3 were also small or invisible in AD mice (Fig. 3D-I).

**Fig. 3.**
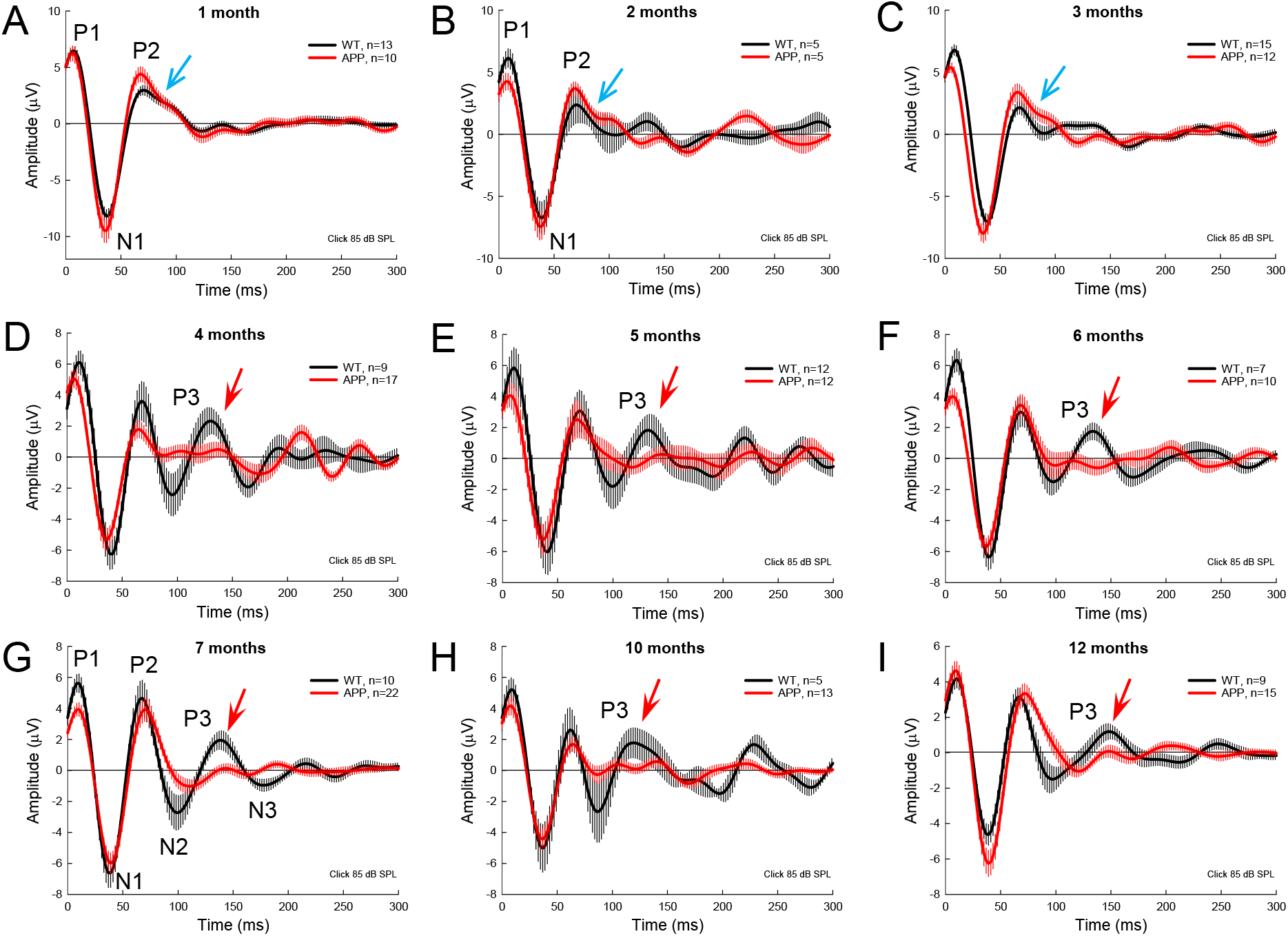
AECP traces recorded from APP/PS1 AD mice at different ages. WT littermates served as control. Red and black traces represent averaged AECP traces recorded from APP/PS1 AD mice and WT mice, respectively. Error bars represent SEM. Mouse age is indicated in each penal. N is the mouse number. Blue arrows indicate a notch at the descending side of peak P2 in AD mice at 1-3 months old. Red arrows indicate the P3 peak clearly visible in WT mice but not in AD mice after 3 months old.

### Changes of AECP in APP/PS1 AD mice with aging

To quantify changes of AECP in AD mice, we measured peaks of AECP at different ages (Fig. 4). Quantitative measurement shows that the early P1 peak in AD mice at 3-7 months old was significantly reduced in comparison with those in WT mice (P=0.014-0.046, t test, two-tail) (Fig. 4A). The amplitudes of later cognitive P3 peak in AD mice were also significantly smaller than WT mice after 3 months old (P=0.049-0.005, t test, two-tail) (Fig. 4E). However, the peak N1 and P2 in AD mice had no significant changes except 4 months old, at which they had significant reduction in comparison with WT mice (P=0.046-0.048, t test, two-tail) (Fig. 4B&C). Later peaks N2 and N3 in AD mice also had no significant reduction in comparison with WT mice except 10-12 months old, at which the N3 peak in AD mice was significantly reduced (P=0.024-0.047, t test, two-tail) (Fig. 4F).

**Fig. 4.**
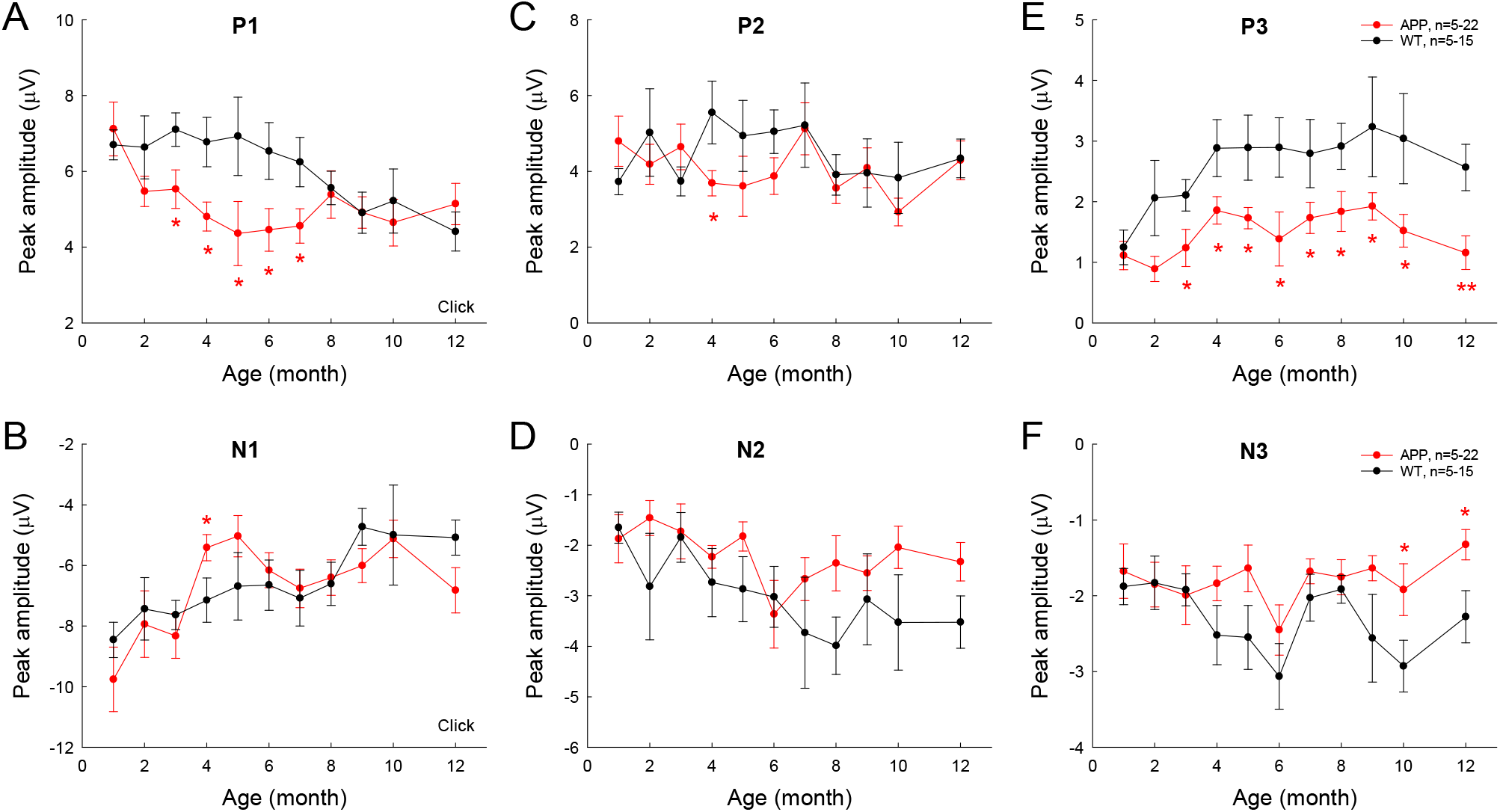
Changes of AECP peaks in APP/PS1 AD mice with aging. WT littermates served as control. Peaks of AECP recorded from each mouse were measured and averaged. Peak name is presented at the top middle place in each panel. N is the mouse number. The amplitudes of peak P3 in AD mice are significantly less than those in WT mice after 3 months old. *: P<0.05, **: P<0.01, t test, two-tail.

The latencies of AECP peaks in AD mice usually had no significant changes or appeared shortening except a few time points (Fig. 5). For example, there was no significant difference in latencies of positive peaks (P1, P2, and P3) between AD and WT mice except 3 months old (Fig. 5A), at which the latency of P1 peak in AD mice was significantly shorter than that in WT mice (P=0.0007, t test, two-tail). For negative peaks, the latency of peak N1 in AD mice appeared shorter than that in WT mice (P=0.0003-0.02, t test, two-tail) at 1, 3, and 6 months old (Fig. 5B). The latency of N3 peak in AD mice also appeared shorter than those in WT mice (P=0.012-0.049, t test, two-tail) at 10-12 months old (Fig. 5F). However, the latency of peak N2 in AD mice appeared longer than that in WT mice (P=0.049, t test, two-tail) at 12 months old (Fig. 5D).

**Fig. 5.**
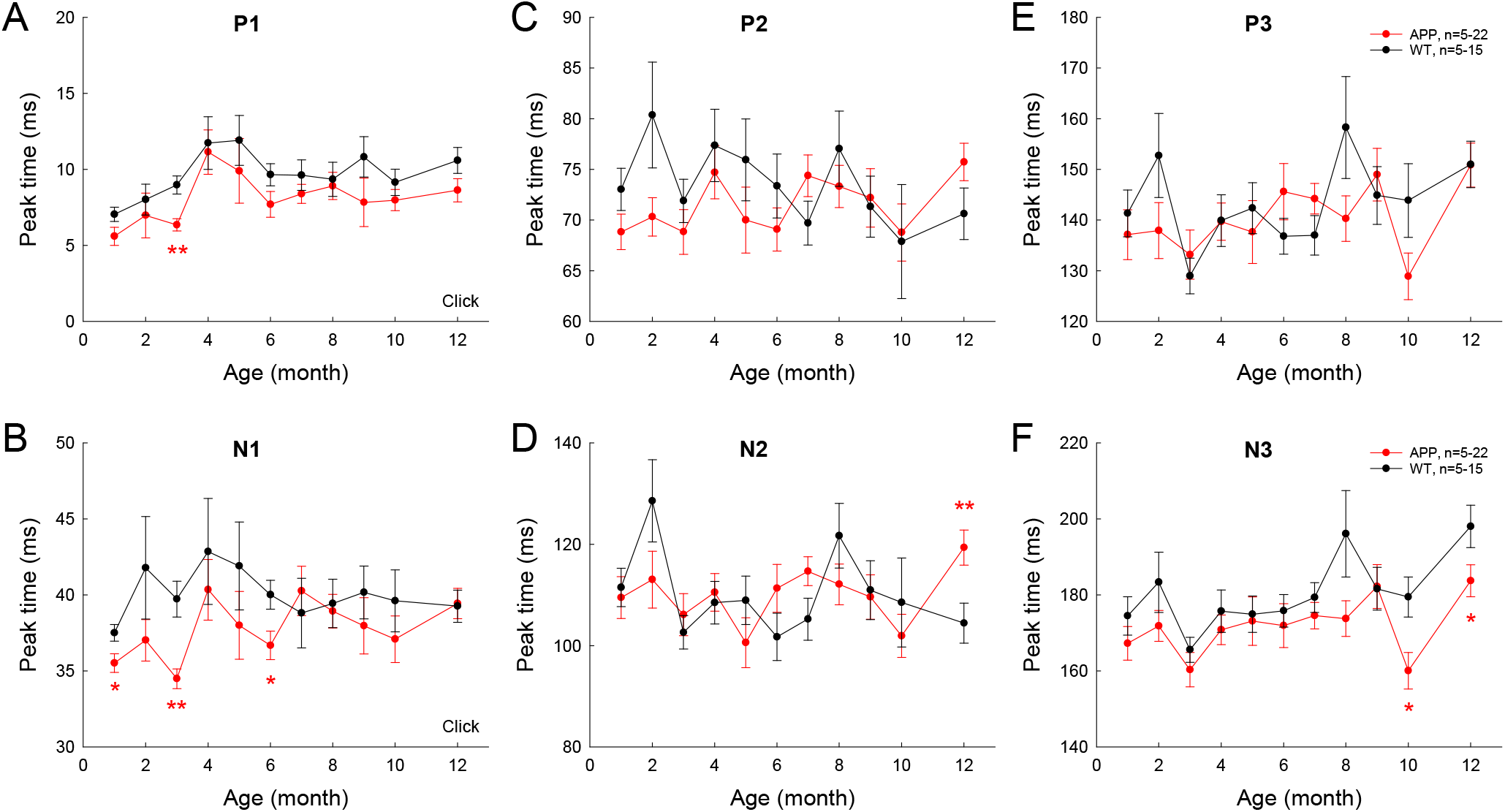
Changes of AECP peak-times in APP/PS1 AD mice with aging. Peak name is presented at the top middle place in each panel. N is the mouse number. The peak-times in AD mice were compared with those in WT littermates at the same age. *: P<0.05, **: P<0.01, t test, two-tail.

### Different changes of AECP in AD mice and CBA/CaJ mice with aging

We further examined AECP changes in AD mice, WT littermate mice, and CBA mice at different ages (Fig. 6). Changes of AECP in WT littermates and CBA mice with normal aging were similar. However, P1 peak in AD mice had significant reduction at 3 and 6 months old (P=0.024-0.035, one-way ANOVA with a Bonferroni correction) (Fig. 6A). Also, P3 peak in AD mice at 3-12 months old was significantly reduced in comparison with that in WT littermates and CBA mice (P=0.0005-0.036, one-way ANOVA with a Bonferroni correction) (Fig. 6E). The later peaks of N2 and N3 in AD mice were also significantly reduced at 12 months old (P=0.004-0.024, one-way ANOVA with a Bonferroni correction) (Fig. 6D&F). However, there were no significant differences in N1 and P2 peaks among AD mice, WT littermates, and CBA mice (Fig. 6B&C).

**Fig. 6.**
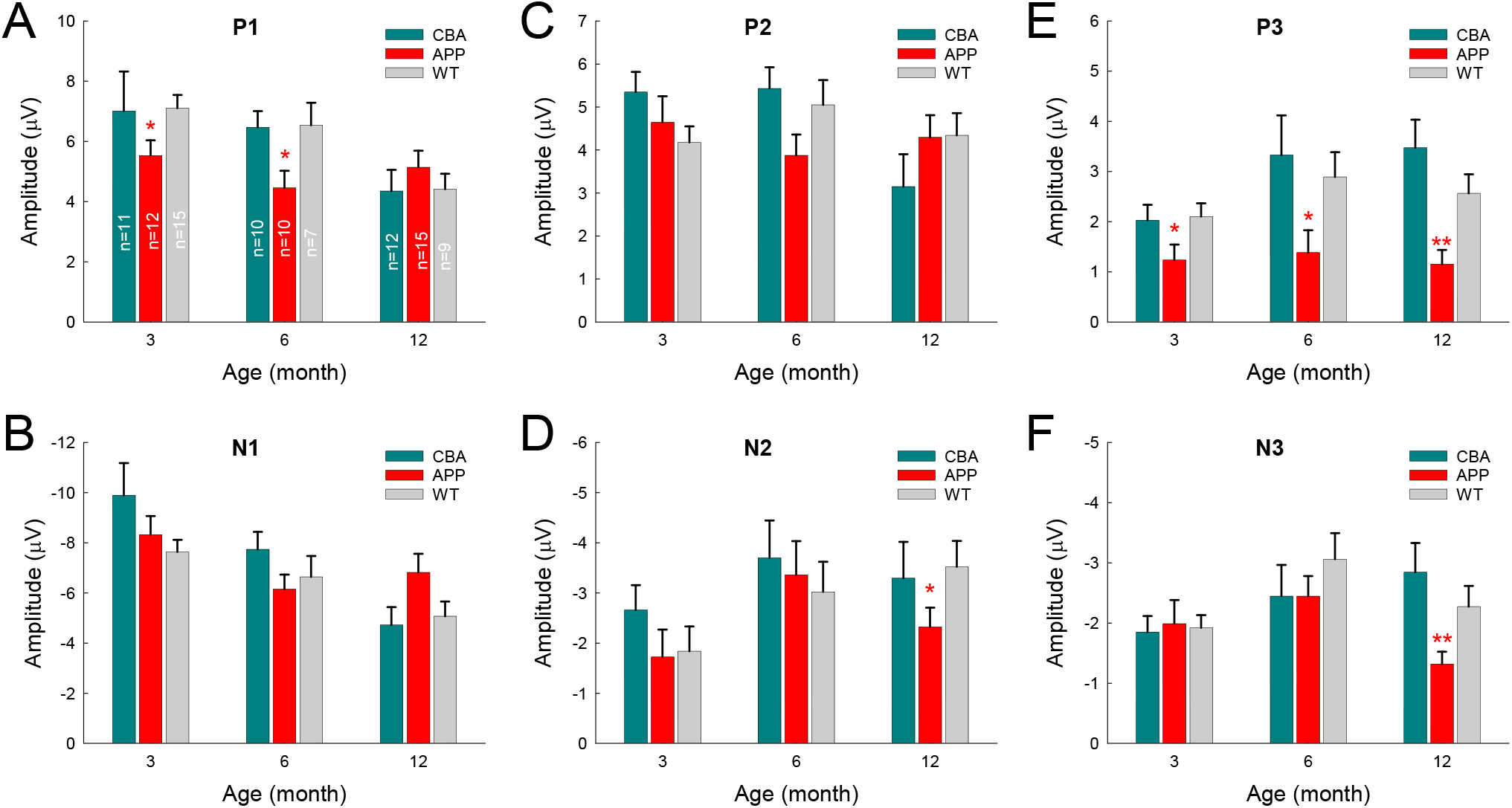
Comparison of AECP changes in CBA/CaJ mice, APP/PS1 AD mice, and WT littermate mice. Peak name is presented at the top middle place in each panel. Animal numbers in each group are presented within the bars in panel A. Panel E shows that peak P3 in the AD mice has significant reduction in comparison with those in CBA and WT littermate mice. *: P<0.05, **: P<0.01, one-way ANOVA with a Bonferroni correction.

In order to assess the aging effect on AECP in AD mice, WT littermates, and CBA mice, the amplitudes of AEACP peaks at different ages were normalized to those at 3 months old in each group. Fig. 7 shows that the early sensory P1, N1, and P2 peaks in CBA mice and WT littermates were decreased as age increased, and had significant reduction at the age of 12 months old in comparison with those at 3 months old (P=0.0003-0.02, t test, two-tail). However, the peak P1, N1, and P2 in AD mice had no significant reduction or change with aging (Fig. 7A-C). For later cognitive peaks, N2, P3, and N3 in CBA mice and WT littermates appeared increased with aging (Fig. 7D-F). In comparison with 3 months old, the increases of N2 in WT littermates at 12 months, P3 peak in CAB mice at 12 months, and N3 peak in WT littermates at 6 months old were significant (P=0.023-0.037, t test, two-tail). However, AECP in AD mice had no significant changes as age increased, except N3 peak at 12 months was significantly decreased in comparison with that at 6 months old (P=0.006, t test, two-tail) (Fig. 7F).

**Fig. 7.**
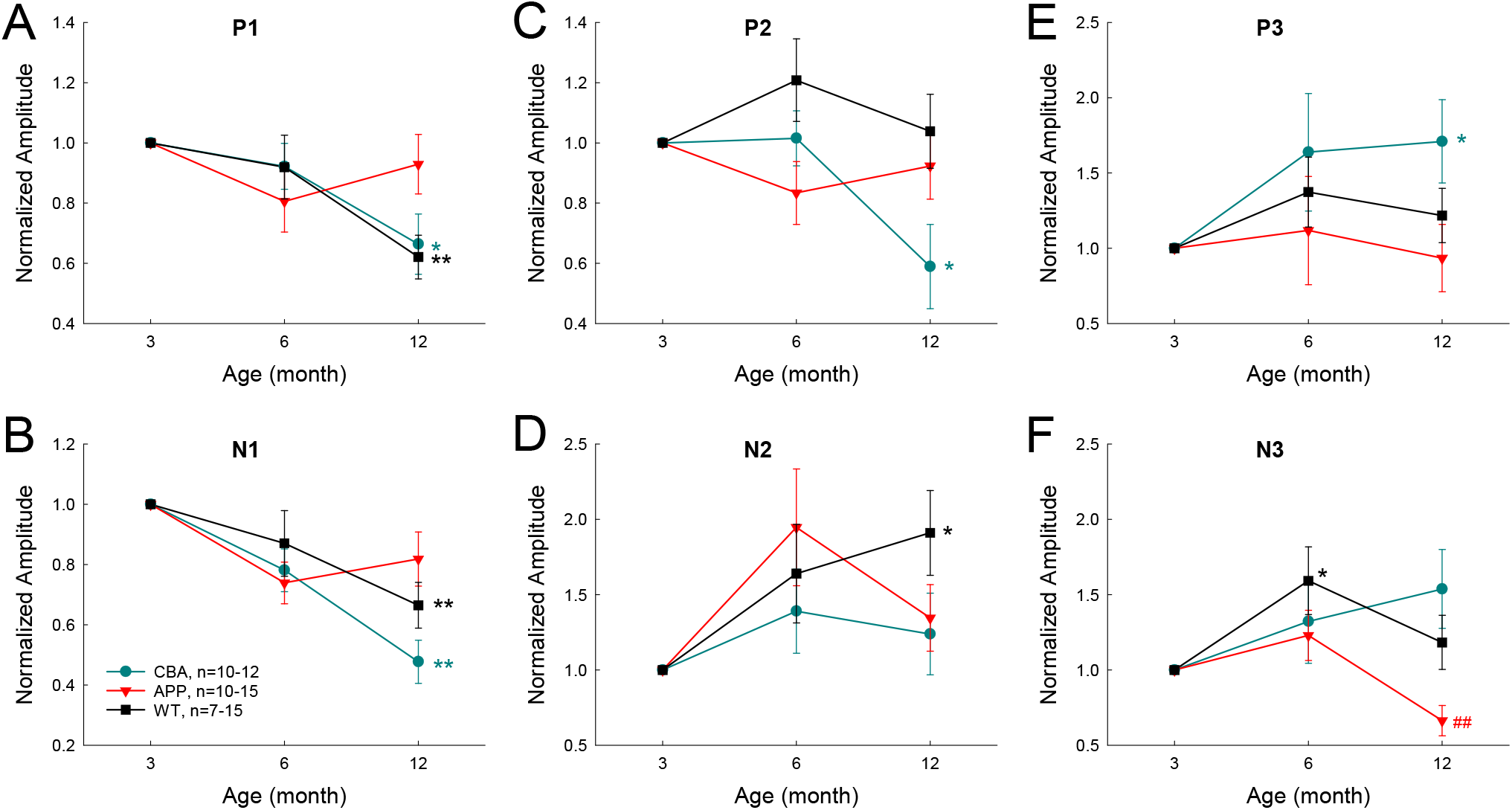
Effects of aging on AECP in CBA/CaJ, APP/PS1 AD, and WT littermate mice. The amplitudes of peaks were normalized to those at 3 months old in each group. Peak name is presented at the top middle place in each panel. N is the animal number. The statistical t test with two-tail was performed by comparing to data at 3 months or data at 6 months in each group. Early AECP peaks (P1, N1, and P2) in CBA mice and WT littermates are decreased as age increased, while later peaks (N2, P3, and N3) are increased or not changed with aging. However, AD mice have no significant changes in AECP peaks with aging except peak N3 at 12 months old, which is decreased. *: P<0.05, **: P<0.01, t test, two-tail, *vs* 3 months; ##: P<0.01, t test, two-tail, *vs* 6 months.

### Expression of amyloid precursor protein (APP) at the auditory cortex in APP/PS1 AD mice

Since APP/PS1 transgenic mice possess both chimeric mouse/human APP (Mo/HuAPP695swe) and mutant human presenilin 1 (PS1-dE9) (Reiserer et al., 2007), we used anti-APP antibody [clone Y188, aa750 to the C-terminus] (#AB32136, Abcam, USA) for immunofluorescent staining. This antibody can also react with gamma secretase fragments and C-terminal fragments. Immunofluorescent staining shows that the positive labeling was visible at the auditory cortex in APP/PS1 AD mice at 2 months old (Fig. 8B&E), consistent with early changes in the AECP recording. However, there was no labeling visible in WT control mice at the same age (Fig. 8C). As age increased, the deposition of labeling in the auditory cortex was increased. At the age of 1 year old, the typical accumulating pattern of plaque deposition was clearly visible at the auditory cortex (Fig. 8A&D).

**Fig. 8.**
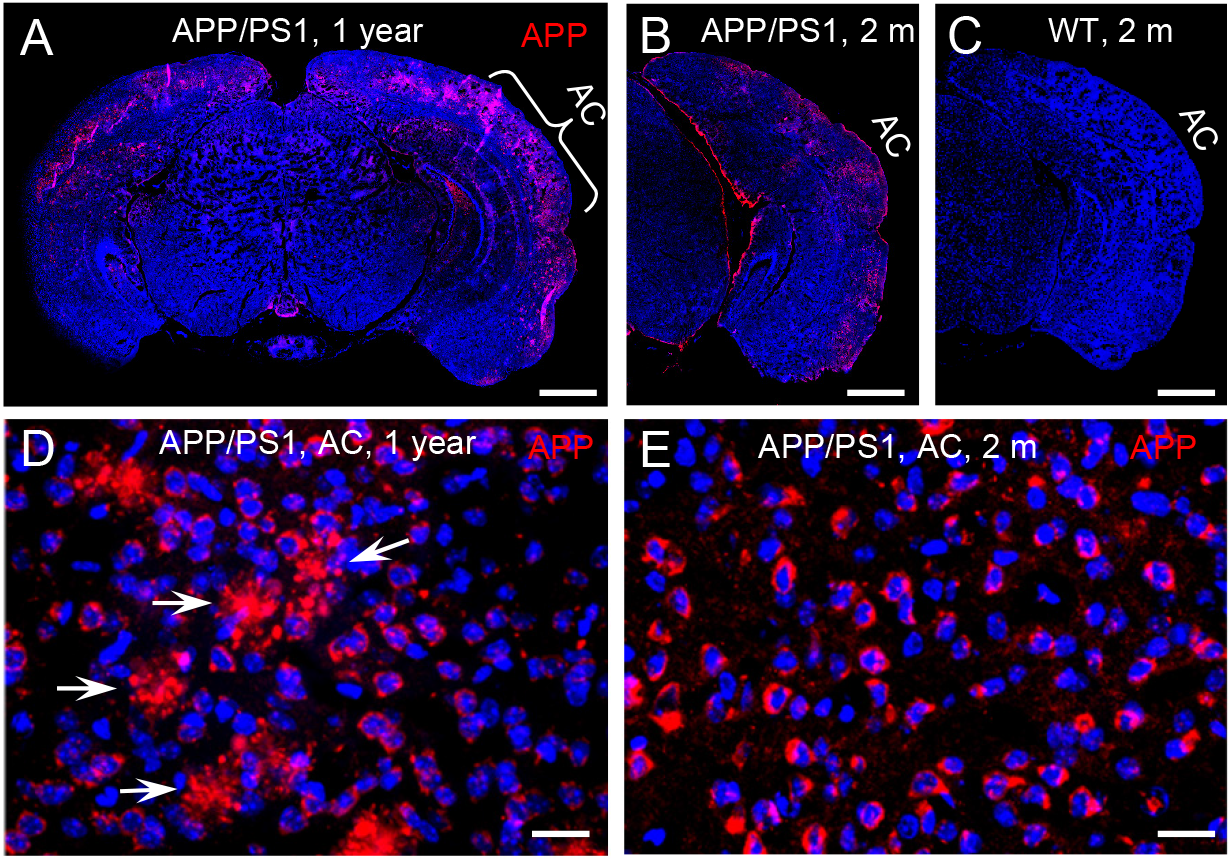
Expression of amyloid precursor protein (APP) in the auditory cortex (AC) in APP/PS1 AD mice. **A-C:** Immunofluorescent staining of brain slices in APP/PS1 AD mice and WT mice at the ages of 2 and 12 months old. **D-E:** High-magnitude images of positive labeling in the AC area. The positive labeling is visible at the AC area in the APP/PS1 AD mouse at 2 months old. White arrows indicate plaque deposits in the AC area at the 12 months old mouse. Scale bar: 100 μm in **A-C**, 10 μm in **D-E**.

## Discussion

In this study, we used the same subdermal needle electrodes and setting as the ABR recording and recorded AECP in mice and characterized changes of AECP in APP/PS1 AD mice and normal aging CBA/CaJ mice. Like the AECP recorded from humans, both early sensory and later cognitive waves were visible in the recorded mouse AECP, which usually presented three waveforms (Fig. 1). The later cognitive peak P3 was not clearly visible until 3 months old (Fig. 1B). As age increased, the early peaks (P1, N1, and P2) in AECP were reduced, while later peaks (i.e. N2, P3 and N3) were increased or had no changes (Figs. 1, 2, and 7). In APP/PS1 AD mice, P1 peak at young ages (<8 months old) appeared smaller than those in WT mice (Figs. 3, 4, and 6). In particular, the later cognitive peak P3 in AD mice was diminished (Figs. 3, 4, and 6). As age increased, there were no significant changes in AECP in AD mice except peak N3 at 1 year old (Fig. 7). Also, there was no significant peak delay in AD mice except peak N2 at 1 year old (Fig. 5). Finally, consistent with early changes of AECP in young ages, intensive labeling for APP was visible in the auditory cortex of AD mice at 2 months old (Fig. 8). These data suggest that AECP could serve as an early, noninvasive biomarker to detect and assess AD development and progression.

In previous studies, various auditory evoked potentials were recorded from animals with implanted electrodes in the brain (Wang et al., 2003; Gurevicius et al., 2013; Kim et al., 2020). However, different from AECP recorded from humans, only early sensory peaks P1 and P2 were visible in those animal recordings; later cognitive peaks were usually missing or invisible even in the recording in the waking animals. In this study, we used subdermal needle electrodes with the same electrode configuration (or setting) as ABR recording and recorded AECP from ketamine-anesthetized mice. The recorded AECP waveforms are like AECP recorded from humans (Golob et al., 2009; Swords et al., 2018); both early sensory and later cognitive peaks were visible in the recording (Figs. 1&3). However, in comparison with the AECP recorded from humans, the AECP in mice appeared only 3 waveforms (Fig. 1). Also, the latencies of P1, P2, and P3 in mice were in the range of 10-15, 65-70, and 110-150 ms, respectively (Fig. 2A&B) and appeared much shorter than humans (Golob et al., 2009; Swords et al., 2018). This may be due to the mouse brain size is small and the neural interactions are less than humans. In addition, P3 in mice was not well-developed until 3 months old (Fig. 1). As mentioned above, later waveforms in AECP are associated with cognitive processing (Sur and Sinha, 2009). Specifically, P3 is dependent on internal thought process and is considered to be mainly generated by the hippocampus where short-term memory is stored. These data suggest that such cognition-associated neural activity in mice may need to take longer time for maturation in the development. These data are also consistent with previous reports that memory-based detection and neural activities are recordable in the anesthetized mice (Chen et al., 2015; Kurkela et al., 2018).

The sensory P1 peak in AD mice appeared smaller than WT mice at young ages (<8 months old) (Figs. 3, 4, and 6). These data are consistent with previous reports that AD or amyloid injection could reduce evoked potentials in the auditory cortex and other brain cortexes in animal models (Wang et al., 2003; Stoiljkovic et al., 2019; Kim et al., 2020; Hidisoglu and Yargicoglu, 2020). These data also indicated that AD could result into functional changes in the auditory cortex at the early AD stage. It has been reported that APP/PS1 AD mice usually appear typical AD phenotypes, such as spatial learning deficit, by age of 6-7 months old (Reiserer et al., 2007; Ordonez-Gutierrez et al., 2015, 2016). We previously reported that hearing loss measured as ABR threshold occurred in APP/PS1 AD mice as early as 2-3 months old; ABR wave IV –V were reduced or diminished (Liu et al., 2020). 5xFAD mice also had early onset of deficits in auditory gap detection at 2 months old (Kaylegian et al., 2019; Weible et al., 2020). These data suggest that AECP recording as well as ABR recording and other hearing function tests could provide important information for early AD development and progression. These data also support the concept that auditory sensory and cognitive cortical potentials in AD persons could be abnormal before AD becomes apparent (Golob et al., 2009).

In this study, consistent with AECP recording, we found that the accumulation of APP in the auditory cortex could be visible in AD mice as early as 2 months old (Fig. 8). As age increased, extracellular Aβ-like plaque deposits were also visible (Fig. 8D). Previous studies demonstrated that the human brainstem and auditory cortex in old AD patients had AD pathology including Aβ plaques, tau protein aggregation, and neural degeneration (Ohm and Braak, 1989; Sinha et al., 1993; Baloyannis et al., 2007, 2009). Our results are consistent with these previous reports and provide further morphological evidence that AD could occur in the auditory system at the early stage.

In comparison with WT littermates, AECP in AD mice usually appeared small (Figs. 3, 4, and 6). This is consistent with previous reports that AD patients had changes in auditory evoked potentials (Buchwald et al., 1989; Irimajiri et al., 2005; Swords et al., 2018). In particular, the later cognitive peak P3 in AD mice was diminished (Figs. 3, 4, and 6), different from normal aging effect (Fig. 7). This is also consistent with a previous report that different from normal aging effect, cognitive N200 and P300 peaks (i.e., N2 and P3 peaks in mice) in AD patients were reduced (Morrison et al., 2018). We further found that as age increased, the early peaks (P1, N1, and P2) in normal aging CBA mice and WT littermates were decreased but the later peaks (N2, P3, and N3) were increased or not changed (Figs. 2 and 7), although the CBA mice have no significant hearing loss as measured by ABR at least until 1 year old (Zheng et al., 1999). Thus, these data implicate that AECP is not only dependent on brainstem input, in particular, the later peaks in AECP, which are mainly generated by the hippocampus associated short-term memory (Sur and Sinha, 2009).

The AECP in AD mice had significant reduction in comparison with WT littermates (Figs. 3, 4, and 6). However, the AECP in AD mice had no significant changes with aging (Fig. 7). This may be due to AECP in AD mice is already reduced and cannot be further reduced as aging increased. In addition, there were no significant delay of AECP peaks in AD mice (Fig. 5), although the latencies of peaks of AECP in normal aging CBA mice were reduced with age increased (Fig. 2B). Actually, the latencies of peak P1 and N1 in AD mice at 1-3 months old (Fig. 5B-D) and the latencies of N3 at 10-12 months old (Fig. 5F) were shorter than those in WT mice. This may arise from the fact that the neural interactions in AD mice may be less than WT mice due to neural degeneration in AD mice.

At present, more and more AD mouse models are available and provide useful animal models to study AD pathological mechanisms, develop therapeutic interventions, and assess the treatment efficiency. However, since lack of reliable objective biomarkers, it is still a big challenge to assess cognitive decline in animals, in particular, at the early AD stage. In this study, we found that later cognitive peak P3 of AECP in AD mice was diminished, different from normal aging effect (Figs. 3, 4, 6, &7). These data suggest that AECP could serve as an objective recording to assess cognitive decline in mice. This will greatly facilitate AD animal studies.

In this study, we reported a new method by using subdermal needle electrodes with the same electrode configuration as ABR recording to record AECP in anesthetized mice. The recorded AECP was similar to AECP recorded from humans, had both early sensory and later cognitive peaks, and showed distinct response patterns in AD mice from normal aging effect (Figs. 1-3, 6, &7). These data suggest that hearing functional tests, including AECP recording, could serve as simple, noninvasive, repeatable biomarkers to assess AD and ADRD development and progression.

## Acknowledgement

This work was supported by NIH R01 DC 017025 and AD supplement DC 017025-01S1 to HBZ.

## Author Contributions

HBZ conceived the general framework of this study. ML, LML, KC and HBZ performed the experiments. ML, LML, and HBZ analyzed data. HBZ wrote the paper. All authors reviewed the manuscript and provided the input.

